# Mental practice modulates functional connectivity between the cerebellum and the primary motor cortex

**DOI:** 10.1101/2021.06.18.448667

**Authors:** Dylan Rannaud Monany, Florent Lebon, William Dupont, Charalambos Papaxanthis

**Author notes:** Corresponding author: Dylan Rannaud Monany, **Email:**. Equal contribution (Florent Lebon, Charalambos Papaxanthis).

## Abstract

Our brain has the extraordinary capacity to improve motor skills through mental practice. Conceptually, this ability is attributed to internal forward models, which are neural networks that can predict the sensory consequences of motor commands. While the cerebellum is considered as a potential locus of internal forward models, evidence for its involvement in mental practice is missing. In our study, we employed single and dual transcranial magnetic stimulation technique to probe the level of corticospinal excitability and of cerebellar-brain inhibition, respectively, before and after a mental practice session or a control session. Motor skills (i.e., accuracy and speed) were measured using a sequential finger tapping-task. Here, we show that mental practice enhances both speed and accuracy. In parallel, the functional connectivity between the cerebellum and the primary motor cortex changes, with less inhibition from the first to the second, expressing the existence of neuroplastic changes within the cerebellum after mental practice. These findings reveal that the corticocerebellar loop is a major neural circuit for skill improvement after mental practice.

## Introduction

A remarkable feature of our brain is its ability to create mental images of past and future events. Part of this mental process is motor imagery, i.e., the internal simulation of body movements without execution (Ruffino et al., 2021). Professional athletes, dancers, and musicians, as well as patients with sensorimotor deficits, use mental practice to improve their motor performance (Schuster et al., 2011, for review). The concept of internal forward models offers the theoretical basis to understand the mental practice process and the associated changes in motor behavior (Kilteni et al., 2018). An internal forward model is a neural network that mimics the causal flow of the physical process by predicting the future sensorimotor state (e.g., position, velocity) given the goal of the movement, the efferent copy of the motor command, and the current state (Wolpert & Flanagan, 2001). Both executed and mental movements seem to share this process. During movement execution, predictions are compared with sensory feedback from the periphery. Any discrepancy constitutes an error signal that can update the internal forward model (i.e., better predictions) and the controller (i.e., better motor commands). During mental movements, such comparison is not possible. However, the goal of the action (e.g., a specific dancing figure) is compared with the prediction from the forward model (i.e., how the dancing figure would be if executed). Any difference between the prediction and the goal acts as an internal error signal that can update and improve the subsequent motor command via a “self-supervised process”.

Neurophysiological and clinical studies consider the cerebellum as a potential locus of internal forward models (Honda et al., 2018; Izawa et al., 2012). Update of motor predictions and motor commands would imply, among others, neuroplastic changes of the neural pathways between the cerebellum and cortical motor areas (Celnik, 2015). Dual coil transcranial magnetic stimulation (TMS) is particularly appropriate to indirectly probe cerebellar adaptations by measuring the influence exerted by the cerebellum onto the primary motor cortex (M1). Specifically, the indirect inhibitory influence of Purkinje cells onto M1 (called cerebellar-brain inhibition, or CBI) diminishes after motor practice (Baarbé et al., 2014). Intriguingly, while behavioral studies confirm the positive effects of mental training in motor learning through forward model predictions, neural evidence supporting them is missing.

## Results & Discussion

Here, we investigated whether improvement in motor performance after mental practice involves neural changes within the cerebellum. We designed an experiment (Fig. 1) in which motor skill in a sequential finger-tapping task and CBI was tested before and after an acute session of mental practice (MP group) or an attentional task (Control group). Movement speed (total number of executed sequences) and accuracy (number of correct sequences) were the markers of motor performance (Walker et al., 2003).

**Figure 1.**
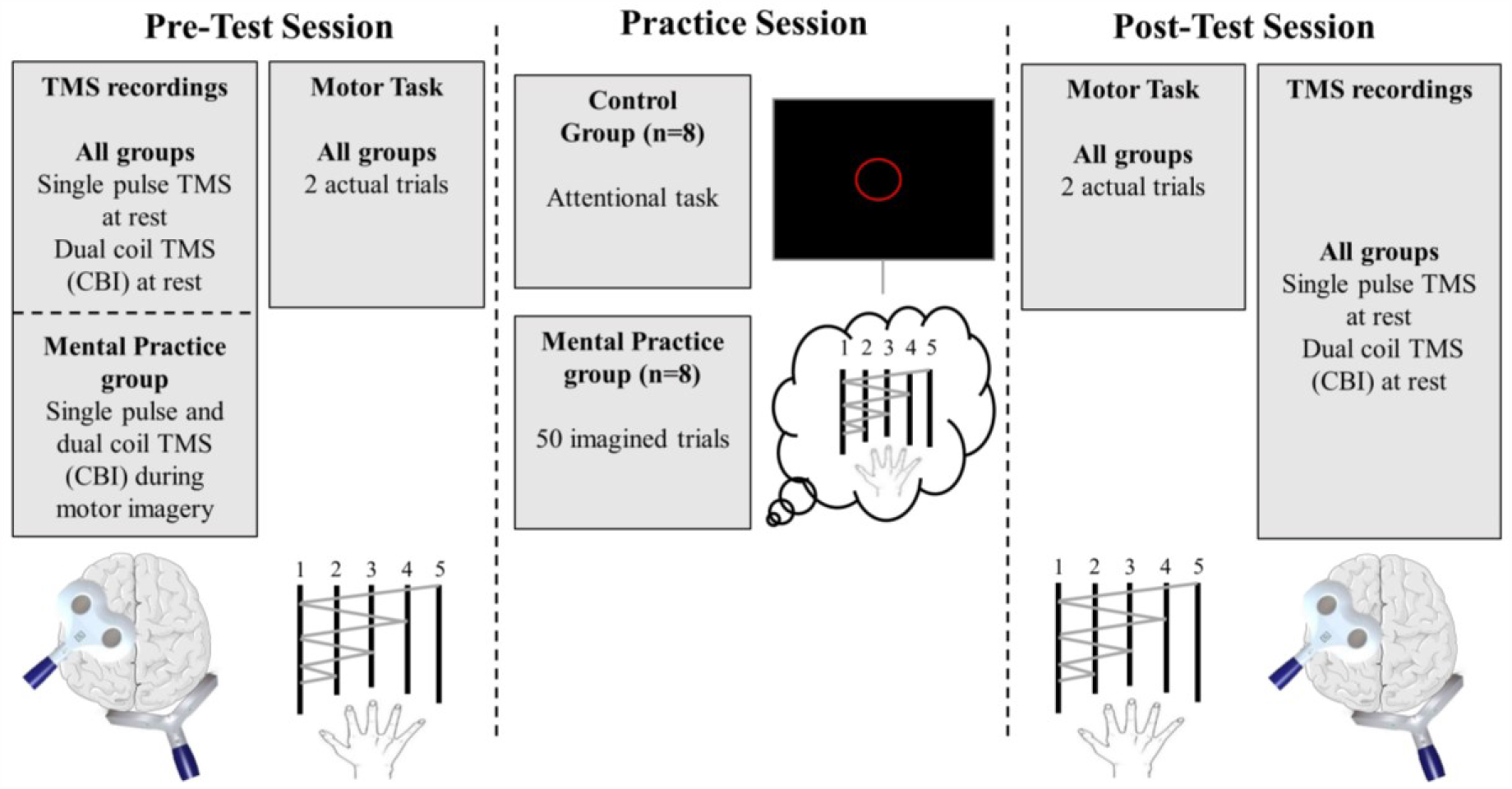
Schematic representation of the experimental procedure. At Pre-Test and Post-Test sessions, both groups executed two trials of the sequential finger-tapping task (order: 1 – 2 – 1 – 3 – 1 – 4 – 1 – 5) with their right hand, as fast and accurately as possible. Each number corresponds to a digit (1= thumb; 5= pinkie). Each trial lasted 10 seconds. Movement speed was defined as the total number of executed sequences per trial, independently of their accuracy. Accuracy was defined as the total number of correct sequences executed per trial. Corticospinal excitability and Cerebellar-Brain Inhibition (CBI) were also assessed in both groups. Neurophysiological measurements were made at rest, i.e., the participants remained quiet without performing any task. Also, CBI was probed during imagined movements for the mental practice group at Pre-Test. During the training session, the Control group performed an attentional task, consisting of counting and memorizing a given number of red circles interspersed within white circles. The mental practice group performed 5 blocks of 10 imagined trials. The duration of both tasks was equivalent.

We found a Group*Time interaction for both movement speed (*F*_1,14_=5.09, *p*=0.04, η_*p*_^2^=0.26) and accuracy (*F*_1,14_=8.79, *p*=0.01, η_*p*_^2^=0.38). Bonferroni post-hoc tests confirmed that the MP group accomplished faster (Fig. 2A) and more accurate (Fig. 2B) the sequential finger-tapping task after training (*Pre-Test vs. Post-Test*; all *p’s*<0.01). The same comparisons were not significant for the Control group (all *p’s*>0.5). It is worth noting that fingers’ muscles remained quiescent during mental practice (*X*^*2*^=7.54, *p*=0.11), excluding any potential influence of muscle activation in skill improvement. Additionally, none of the groups showed mental fatigue after training (see complementary results, section A).

**Figure 2.**
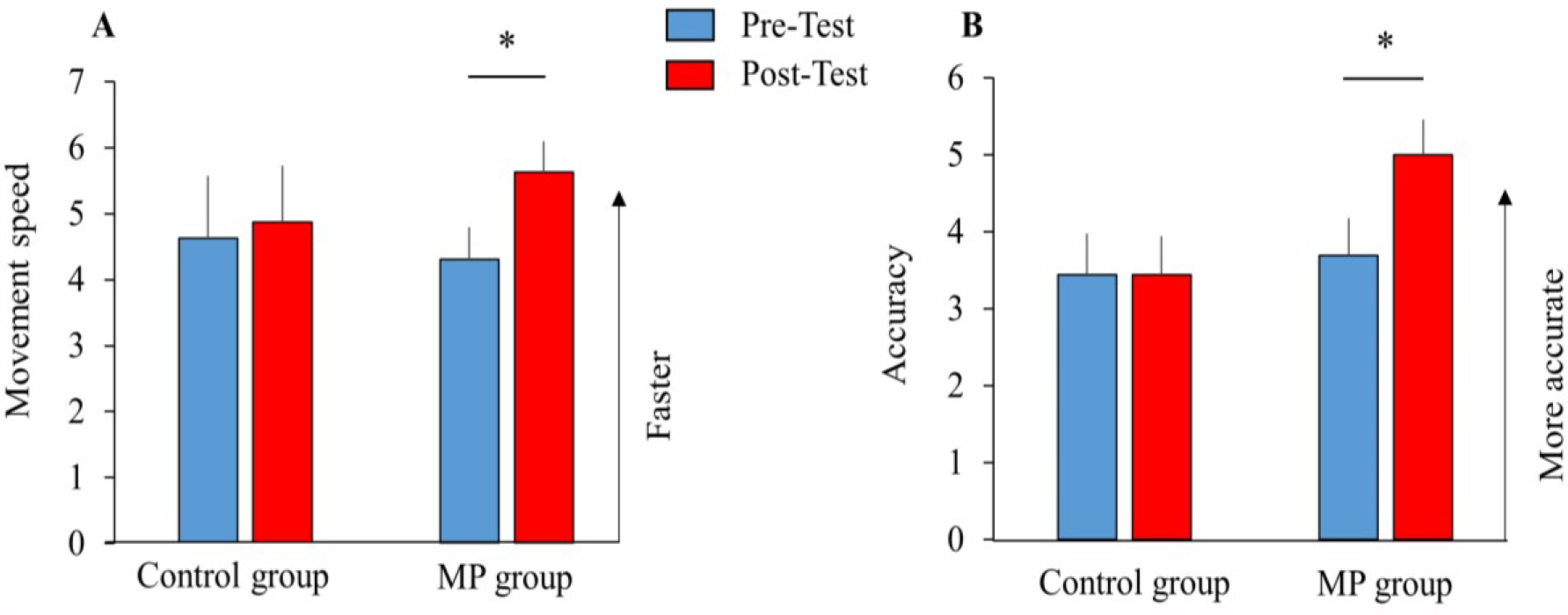
Mean ± SE (standard errors) of movement speed (i.e., the total number of executed sequences, **A**) and accuracy (i.e., the number of correct sequences, **B**) for both groups at Pre-Test and Post-Test. Both parameters significantly increased following mental practice (MP), but not after the attentional task (Control group). ^*^: *p*<0.05.

To probe neural changes within the cerebellum, we measured CBI between the right cerebellum and left M1 using dual coil TMS. Note that before testing CBI, we first verified that the sole stimulation at the cervicomedullary junction did not elicit descending volleys into the spinal cord (see complementary results, section B) and that the TMS parameters were similar between the two groups (see complementary results, section C). Thereafter, we found a modulation of CBI following mental practice, attested by the significant Group*Time interaction (*F*_1,14_=6.38, *p*=0.024, *ηp2*=0.31, Fig. 3). Interestingly, Bonferroni post-hoc tests revealed that CBI was no longer present after practice for the MP group (- 8.91 ±3.01% at Pre-Test vs. -0.28 ±6.04% at Post-Test, *p*=0.024), whereas it was still observed after the attentional task for the Control group (−12.11 ±11.53% at Pre-Test vs. - 12.48 ±11.79% at Post-Test, *p*=0.94).

**Figure 3.**
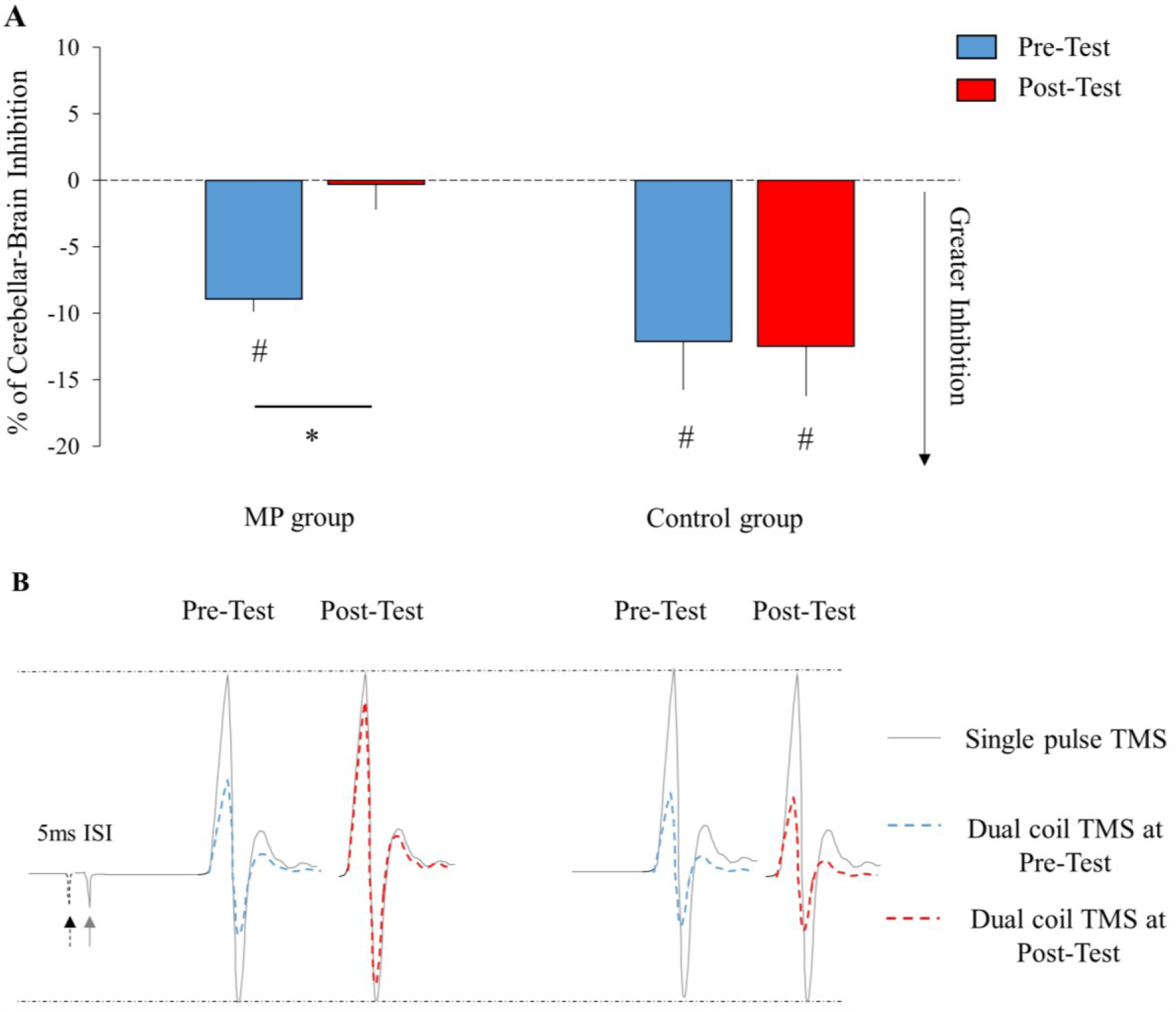
**A**. Mean ± SE (standard error) of the percentage of Cerebellar-Brain Inhibition (CBI) at Pre-Test and Post-Test sessions for both groups. The main finding is a disinhibition of CBI following mental practice (MP). We used one-sample t-test against 0 to ensure the presence of CBI for the Control group at Pre-Test: (−12.11 ±11.53%, *t*(7)=2.97, *p*=0.02, *Cohen’s d*=1.05) and Post-Test (−12.48 ±11.79, *t*(7)=2.99, *p*=0.02, *Cohen’s d*=1.06). For the MP group, CBI values were different from 0 at Pre-Test (−8.91 ±3.01%, *t*(7)=8.38, *p*<0.01, *Cohen’s d*=2.96), but not at Post-Test, indicating no-inhibition (−0.28 ±6.04%, *p*=0.9). ^*^: *p*<0.05 (comparison between Pre- and Post-Test); #: *p*<0.05 (comparison to 0). **B**. Qualitative representation of CBI. Single-pulse transcranial magnetic stimulation (TMS) over M1 elicits a motor-evoked potential (MEP) in the target muscle (grey lines). When conditioning M1 with cerebellar stimulation, the MEP amplitude reduces, reflecting CBI (Pre-Test, blue dotted lines). While the conditioned MEP remains reduced for the Control group, it increases following MP showing a disinhibition mechanism (Post-Test, red dotted lines). ISI: inter-stimulation interval.

It is worth mentioning that the disinhibition was due to plastic changes that occurred within the cerebellum. Indeed, corticospinal excitability at rest (single-pulse TMS condition) was not modulated (see complementary results, section D), excluding neural adaptations between M1 and the target muscle after an acute session of mental practice. In addition, the disinhibition observed at rest after practice can be directly attributed to CBI modulation occurring during imagined movements. Indeed, we found a significant reduction of CBI while imagining compared to rest at Pre-Test (−2.62 ±6.64% and -8.91 ±3.01%; *t*(14)=2.44, respectively, *p*=0.029, *Hedges’s g*=0.97).

We provided evidence that an acute session of mental practice improves motor performance and induces neural adaptations within the cerebellum (i.e., reduction of CBI). These results reinforce the premise regarding the positive contribution of internal movement prediction during consecutive imagined actions (Ruffino et al., 2021, Kilteni et al., 2018). The state predictions (e.g., position and speed) generated by the forward model during mental practice are appropriate to elicit functional adaptations within the corticocerebellar loop and thus improve motor skills (i.e., faster and more accurate movements). Clinical studies support this statement, showing that cerebellar lesions affect the motor imagery process of complex motor sequences and alter the update of internal movement predictions (Battaglia et al., 2006; Synofzik et al., 2008; Saunier et al., 2021). The current neurophysiological hypothesis to explain the reduction of CBI is the long-term depression-like plasticity phenomenon in inhibitory GABAergic Purkinje fibers that indirectly project to the contralateral M1 (Ishikawa et al., 2016). Briefly, the reduction of the inhibitory output from the GABAergic Purkinje cells to the deep cerebellar nuclei would release the excitatory activity of these nuclei onto M1. This mechanism has been well established in motor adaptation and motor skill learning studies (Galea et al., 2011) and seems to be also engaged in mental practice. Our results corroborate and extend those of previous studies in skill learning (Spampinato and Celnik., 2017), which showed a significant reduction of CBI immediately after an acute session of physical training. Despite the absence of sensorimotor feedback during mental practice, the cerebellum is at play as part of the internal forward model to predict the sensory consequences of the imagined action (Kilteni et al., 2018), and to adapt motor commands via a putative self-supervised process. In conclusion, the current findings suggest the importance of the corticocerebellar loop in motor learning through mental practice, which can be used in isolation or in addition to actual practice to improve motor performance in healthy individuals or patients with motor deficits.

## Materials and Methods

### Participants

We first conducted a power analysis on G*power (Faul et al., 2007) to determine the required sample size to reach a power of at least 0.8, considering two groups and large effect size (partial eta^2^ = 0.2), extrapolated from previous similar studies (Zabihhosseinian et al., 2020). The results of the analysis gave eight (n=8) participants for each group. Therefore, sixteen healthy right-handed volunteers participated in the study (3 women, mean age: 23.12 years old, range: 20 - 26). All were screened for contraindications to TMS by a medical doctor and had normal or corrected vision. The participants were randomly assigned to the mental practice group (MP group, n=8, mean age: 23, range 20 - 26) and the Control group (n=8, mean age: 23.25, range 20 - 25). The study was approved by the CPP SOOM III ethics committee (ClinicalTrials.gov Identifier: NCT03334526) and complied with the standards set by the Declaration of Helsinki.

## Experimental protocol

### Behavioral task

The motor task was a computerized version of the sequential finger-tapping task (Karni et al., 1998; Debarnot et al., 2009). The participants were comfortably seated in front of a customized keyboard and performed a sequence of eight movements using their right fingers in the following order: 1-2-1-3-1-4-1-5 (1: thumb, 2: index finger, 3: middle finger, 4: ring finger, 5: pinky), as fast and accurately as possible (Figure 1). At Pre-Test and Post-Test sessions, the participants performed two trials of 10s each. Between the test sessions, the MP group imagined the same sequence during five blocks of ten trials (total number of trials = 50). Each trial lasted 10 seconds with 10-second rest. The participants of the MP group placed their right hand on the keyboard, and imagined the motor sequence with the following instructions: “*try to imagine yourself performing the motor task as fast and accurately as possible, by feeling your fingers moving as if you were moving it*”. Electromyographic (EMG) activity was recorded during mental practice to ensure the absence of muscular activity. The Control group performed a visual recognition task, during which the participants counted and memorized the number of red circles interspersed within white circles. The number of red circles varied between blocks to avoid habituation effects. The participants reported the number of red circles they memorized after each block to confirm that they were focused on the task. The duration of this task matched the duration of mental practice. For both groups, we assessed the mental fatigue before and after the tasks, using a 10-cm visual analog scale (0 cm: ‘no fatigue’, 10 cm ‘maximal fatigue’).

### Transcranial Magnetic Stimulation

We assessed the level of corticospinal excitability with single-pulse transcranial magnetic stimulation (TMS) and the amount of cerebellar-brain inhibition (CBI) with dual-coil TMS. Single-pulse and dual-coil TMS were applied using monophasic BiStim^2^ stimulators (*The Magstim Co*., *Whitland, UK*). Motor-evoked potentials (MEPs) were recorded in the right Abductor Pollicis Brevis (APB) muscle.

#### TMS over M1 (Test Stimulus, TS)

Single-pulse stimulations were applied with a 70-mm figure-of-eight coil place over the left M1 in a posterior position at 45° from the sagittal plane to induce a postero-anterior current flow. We identified the hotspot as the site eliciting the highest and most consistent MEPs amplitude in APB. We determined the rest motor threshold (rMT) as the lowest stimulator output that elicited 4 of 8 MEPs with peak-to-peak amplitude equal to or greater than 0.05 mV. Then, we determined the intensity to evoke MEP_max_ at rest, i.e., the individual highest peak-to-peak MEP amplitude. For the rest of the experiment, we considered MEP_target_ as half of MEP_max_ amplitude.

#### TMS over the cerebellum (Conditioning Stimulus, CS)

We used a double-cone coil to stimulate deep cerebellar structures. The double-cone coil was positioned over the cerebellum on the horizontal line between the inion and the external auditory meatus at 2 cm right to the inion. The stimulation intensity was fixed at 150% of individual M1 rMT. Although the cerebellar stimulation intensity in the current experiment was lower than that used in previous studies (Bunday et al., 2014), we delivered five cerebellar stimulations alone at rest at Pre-Test to verify the absence of descending volleys in EMG traces.

#### Dual-coil stimulations (CS and TS)

Dual-coil stimulations were applied through the combination of the double-cone coil TMS (CS) over the right cerebellum and a 70-mm figure-of-eight coil over left M1 (TS, see above). The inter-stimulation interval between CS and TS was set to 5ms to ensure the activation of cerebellar-M1 inhibitory pathways (Fisher et al., 2009).

### Adaptive threshold-hunting technique

This method has been recently developed to overcome the potential limitations of conventional paired-pulse TMS protocols, such as large variability in MEP amplitude and a ‘‘floor/ceiling effect” when the observed inhibition leads to complete MEP suppression (Cirillo and Byblow, 2016; Van der Bos et al., 2018). The hunting-threshold was defined as the TS intensity (expressed in percentage of the maximal stimulator output - %MSO) required to elicit the MEP_target_ (i.e., half of MEP_max,_ see above) in the relaxed APB. To do so, we used an available online freeware (TMS Motor Threshold Assessment Tool, MTAT 2.0), based on a maximum-likelihood Parameter Estimation by Sequential Testing. Following previous studies, the software has been parameterized with “assessment without *a priori* information (Cirillo et al., 2018; Neige et al., 2020). Once the amplitude of MEP_target_ and the intensity of CS (150% rMT) were set, we determined the TS intensity (%MSO) required to maintain the MEP_target_ amplitude across the experimental conditions. Therefore, the TS intensity was the dependent variable of the experiment.

Twenty stimulations were delivered for the Single pulse TMS condition (SingleRest) and twenty stimulations for the dual coil TMS condition (DualRest) at rest in the Pre-test and Post-Test sessions. In addition, the same number of stimulations was derived during single pulse TMS (SingleMI) and dual coil TMS (DualMI) during motor imagery for the MP group only at Pre-Test. The order of the stimulation conditions was randomized across participants.

### Motor imagery questionnaire

Participants of the MP group were asked to complete the French version of the Motor Imagery Questionnaire to assess self-estimation of their motor imagery vividness (Loison et al., 2011). For this questionnaire, the minimum score is 8 (low imagery vividness) and the maximum one is 56 (high imagery vividness). In the current study, the average score of the MP group participants (43± 5.29) suggests good motor imagery vividness.

### Data recording and analysis

#### Behavioral parameters

Two motor parameters were assessed: movement speed and accuracy. These parameters were extracted for each trial and averaged for Pre-Test and Post-Test. Movement speed was defined as the total number of executed sequences per trial, independently of their accuracy. Accuracy was defined as the total number of correct sequences executed per trial.

#### Neurophysiological parameters

Values were quantified as %MSO for the four TMS recordings conditions (i.e., SingleRest, SingleMI, DualRest, and DualMI). To investigate variations of Cerebellar-Brain Inhibition at rest and during MI in dual-coil TMS conditions, we used the following formula:

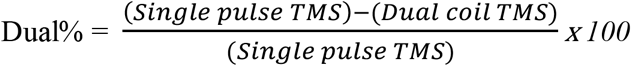

where positive values indicate facilitation and negative values indicate inhibition. DualRest and DualMI were normalized according to SingleRest and SingleMI, respectively.

#### Electromyographic recording and analysis

Electromyographic (EMG) recordings of the right APB muscle were made with surface Ag/AgCl electrodes in a belly-tendon montage. A ground electrode was placed on the styloid process of the ulna. The EMG signals were amplified and band-pass filtered (10–1000 Hz, Biopac Systems Inc.) and digitized at a sampling rate of 2000 Hz for offline analysis. Background EMG was monitored for the 100 ms preceding every TMS pulse to ensure a complete muscle relaxation throughout the experiments, using the following formula:

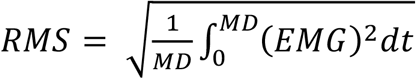

## Statistical analysis

Statistical analysis was performed using STATISTICA (*13*.*0 version; Stat-Soft, Tulsa, OK*). Normality was checked before inferential analysis using Shapiro Wilk tests. Cohen’s d and Hedges’s g (one-sample and paired sample t-tests, respectively), and partial eta squared (general linear model) were reported to provide information on effect sizes. P-values were adjusted accordingly using the Bonferroni method when several tests were performed on the same variable. The threshold of statistical significance was set to α= 0.05. First, movement speed and accuracy were analyzed using a GLM analysis with Time (Pre-Test vs. Post-Test) as a within-subject factor and Group (Control vs. MP) as a between-subject factor. Bonferroni post-hoc comparisons were performed in case of significant Group*Time interaction. A Friedman’s ANOVA (normality was violated) was performed to compare the EMG activity of APB at each imagined block with recording at rest, to ensure that muscles remained silent during MP.

Then, DualRest was analyzed using a GLM analysis with Time (Pre-Test vs. Post-Test) as a within-subject factor and Group (Control vs. MP) as a between-subject factor. Bonferroni post-hoc comparisons were performed in case of significant Group*Time interaction. One-sample t-tests versus 0 were used to test if dual coil TMS induced significant amount of CBI at Pre-Test and Post-Test for Control and MP groups. Also, to test whether the cerebellum is involved during motor imagery before practice (Tanaka et al., 2018), we performed a one-tailed paired-sample t-test opposing DualRest and DualMI for the MP group at Pre-Test.

## Complementary results

### A. Mental fatigue

We used one-sample t-tests against the reference 0 to ensure mental fatigue did not increase after mental practice or attentional task, considering the difference between Post-Test and Pre-Test for each groups (Rozand et al., 2016). There were no statistical differences for the MP group (Pre-Test: 3.37 ±1.45 cm, Post-Test: 3.76 ±1.45 cm, difference: +0.39 ±0.5 cm, *t*(7)=2.20, *p*=0.12) or the Control group (Pre-Test: 2.26 ±1.28 cm, Post-Test: 2.60 ±1.20 cm, difference: +0.34 ±0.41 cm, *t*(7)=2.33, *p*=0.1). Mental fatigue did not significantly increase after mental practice or the attentional task.

### B. Cervicomedullar output

We used one-tailed one-sample t-tests against the reference value 0.05 to ensure that the amplitude of raw EMG following cerebellar stimulations alone remained significantly lower than the rest motor threshold (0.05 mV). We found that EMG traces remained below the rest motor threshold for the Control group (0.02 ±0.02 mV; *t*(7)=-3.45, *p*=0.01, *Cohen’s d*= -1.22) and the MP group (0.01 ±0.01 mV; *t*(7)=-7.15, *p*<0.01, *Cohen’s d*= -2.53), suggesting that cerebellar stimulations alone did not induce descending volleys at the cervicomedullary junction.

### C. Rest motor threshold, MEP_target_ amplitude and cerebellar stimulation intensity

We used independent t-tests to ensure that Control and MP groups were not statistically different regarding the rest motor threshold (rMT), the MEP_target_ amplitude, and the cerebellar stimulation (CS) intensity. The rMT (MP group: 43.12 ±7.58 %MSO, Control group: 44.75 ±7.05 %MSO), the MEP_target_ amplitude (MP group: 0.47 ±0.16 mV, Control group: 0.60 ±0.31 mV), and the CS intensity (MP group: 66.75 ±9.75 %MSO, Control group: 67.12 ± 10.69 %MSO) were not different between groups (all *p’s*>0.1).

### D. Corticospinal excitability (Single pulse TMS)

We used paired sample t-tests to ensure that single-pulse TMS intensities at rest remained stable at Post-Test when compared to those at Pre-Test for both groups (MP group, Pre-Test: 52.97 ±10.47 %MSO; Post-Test: 54.82 ±11.62 %MSO, *t*(7)=0.79, *p*=0.46) and Control group, Pre-Test: 55.65 ± 9.72 %MSO; Post-Test: 56.50 ±10.60 %MSO, *t*(7)=0.56, *p*=0.59). This result confirms that an acute session of MP did not significantly increase corticospinal excitability.

## Author Contributions

DRM, CP, and FL designed the experiment, DRM and WD conducted the experiment, DRM analyzed data, DRM prepared figures, DRM, CP and FL wrote the manuscript, WD, CP and FL provided feedback on the manuscript. All authors read and approved the current version of the manuscript.

## References

Baarbé J, Yielder P, Daligadu J, Behbahani H, Haavik H, Murphy BA (2014) A novel protocol to investigate motor training-induced plasticity and sensorimotor integration in the cerebellum and motor cortex Journal of Neurophysiology 111:4, 715–721. https://doi.org/10.1152/jn.00661.2013

Battaglia F, Quartarone A, Ghilardi AMF, Dattola R, Bagnato S, Rizzo V, Morgante L, Girlanda P (2006) Unilateral cerebellar stroke disrupts movement preparation and motor imagery Clinical Neurophysiology 117, 1009–1016. https://doi.org/10.1016/j.clinph.2006.01.008

Bunday KL, Tazoe T, Rothwell JC, Perez MA (2014) Subcortical Control of Precision Grip after Human Spinal Cord Injury Journal of Neuroscience 34:21, 7341–7350.

Celnik P (2015) Understanding and modulating motor learning with Cerebellar stimulation Cerebellum 14:2, 171–174. https://doi.org/10.1523/JNEUROSCI.0390-14.2014

Cirillo J, Byblow WD (2016) Threshold tracking primary motor cortex inhibition: The influence of current direction. Experimental Brain Research 233, 679–689. https://doi.org/10.1111/ejn.13369

Cirillo J, Semmler JG, Mooney RA, Byblow WD (2018) Conventional or threshold-hunting TMS? A tale of two SICIs Brain Stimulation 11, 1296–1305. https://doi.org/10.1016/j.brs.2018.07.047

Debarnot U, Creveaux T, Collet C, Doyon J, Guillot A (2009) Sleep Contribution to Motor Memory Consolidation: A Motor Imagery Study Sleep 32:12, 1559–1565. https://doi.org/10.1093/sleep/32.12.1559

Faul F, Erdfelder E, Lang AG, Buchner A (2007) G*Power: A flexible statistical power analysis program for the social, behavioral, and biomedical sciences Behavioral Research Methods 39, 175–193. https://doi.org/10.3758/bf03193146

Fisher KM, Lai HM, Baker MR, Baker SN (2009) Corticospinal activation confounds cerebellar effects of posterior fossa stimuli Clinical Neurophysiology 120:12, 2109–2113. https://doi.org/10.1016/j.clinph.2009.08.021

Galea JM, Vazquez A, Pasricha N, Orban de Xivry JJ, Celnik P (2011) Dissociating the Roles of the Cerebellum and Motor Cortex during Adaptive Learning: The Motor Cortex Retains What the Cerebellum Learns Cerebral Cortex 21: 8, 1761–1770. https://doi.org/10.1093/cercor/bhq246

Honda T, Nagao S, Hashimoto Y, Ishikawa K, Yokota T, Mizusawa H, Ito M (2018) Tandem internal models execute motor learning in the cerebellum Proceedings of the National Academy of Sciences of the United States of America 115:28, 7428–7433. https://doi.org/10.1073/pnas.1716489115

Ishikawa T, Tomatsu S, Izawa J, Kakei S (2016) The cerebro-cerebellum: Could it be loci of forward models? Neuroscience Research. 104, 72–79. https://doi.org/10.1016/j.neures.2015.12.003

Izawa J, Criscimagna-Hemminger SE, Shadmehr R (2012) Cerebellar Contributions to Reach Adaptation and Learning Sensory Consequences of Action Journal of Neuroscience 32:12, 4230–4239. https://doi.org/10.1523/JNEUROSCI.6353-11.2012

Karni A, Meyer G, Rey-Hipolito CH, Jezzard P, Adams MM, Turner R, Ungerleider LG (1998) The acquisition of skilled motor performance: Fast and slow experience-driven changes in primary motor cortex Proceedings of the National Academy of Sciences of the United States of America 95:3, 861-868 (1998). https://doi.org/10.1073/pnas.95.3.861

Kilteni K, Anderson BJ, Jouborg C, Ehrsson HH (2018) Motor imagery involves predicting the sensory consequences of the imagined movement Nature Communications 9:1, 1617. https://doi.org/10.1038/s41467-018-03989-0

Loison B, Moussaddaq AS, Cormier J, Richard I, Ferrapie AL, Ramond-Roquin Dinomais M (2013) Translation and validation of the French Movement Imagery Questionnaire – Revised Second Version (MIQ-RS) Annals of Physical and Rehabilitation Medecine 56:3, 157–173. https://doi.org/10.1016/j.rehab.2013.01.001

Neige C, Rannaud Monany D, Stinear CM, Byblow WD, Papaxanthis C, Lebon F (2020) Unravelling the Modulation of Intracortical Inhibition During Motor Imagery: An Adaptive Threshold-Hunting Study Neuroscience 434, 102–110. https://doi.org/10.1016/j.neuroscience.2020.03.038

Rozand V, Lebon F, Stapley PJ, Papaxanthis C, Lepers R (2015) A prolonged motor imagery session alter imagined and actual movement durations: Potential implications for neurorehabilitation Behavioral Brain Research 297, 67–75. https://doi.org/10.1016/j.bbr.2015.09.036

Ruffino C, Truong C, Dupont W, Bouguila F, Michel C, Lebon F, Papaxanthis C (2021) Acquisition and consolidation processes following motor imagery practice Scientific Reports 11:2295. https://doi.org/10.1038/s41598-021-81994-y

Saunier G, Fontana AP, De Oliveira JM, Py MO, Pozzo T, Vargas CD (2021) Cerebellar damage affects the inference of human motion. Neurocase 1–9. https://doi.org/10.1080/13554794.2021.1905853

Schuster C, Hilfiker R, Amft O, Scheidhauer A, Andrews B, Butler J, Kischka U, Ettlin T (2011) Best practice for motor imagery: a systematic literature review on motor imagery training elements in five different disciplines BMC Med. 9:75. https://doi.org/10.1186/1741-7015-9-75

Spampinato D, Celnik P (2017) Temporal dynamics of cerebellar and motor cortex physiological processes during motor skill learning. Scientific Reports 7, 40715. https://doi.org/10.1038/srep40715

Synofzik MA., Lindner AP. Their P (2008) The Cerebellum Updates Predictions about the Visual Consequences of One’s Behavior Current Biology 18, 814–818. https://doi.org/10.1016/j.cub.2008.04.071

Tanaka H, Matsugi A, Okada Y (2018) The effects of imaginary voluntary muscle contraction and relaxation on cerebellar brain inhibition Neuroscience Research 133, 15–20. https://doi.org/10.1016/j.neures.2017.11.004

Van der Bos MAJ, Menon P, Howells J, Geevasinga N, Kiernan MC, Vucic S (2018) Physiological processes underlying short interval intracortical facilitation in the human motor cortex Frontiers in Neuroscience 12, 1–11. https://doi.org/10.3389/fnins.2018.00240

Walker MP, Brakefield T, Seidman J, Morgan A, Hobson JA, Stickgold R (2003). Sleep and the Time Course of Motor Skill Learning Learning and Memory 10:4, 275–284. https://doi.org/10.1101/lm.58503

Wolpert DJ., Flanagan J (2001) Motor prediction Current Biology 11:18, 729–732. https://doi.org/10.1016/s0960-9822(01)00432-8

Zabihhosseinian M, Yielder P, Berkers V, Ambalavanar U, M. Holmes MB. Murphy B (2020) Neck muscle fatigue impacts plasticity and sensorimotor integration in cerebellum and motor cortex in response to novel motor skill acquisition Journal of Neurophysiology 124:3, 844–855. https://doi.org/10.1152/jn.00437.2020

